# Long-term trends in the contribution of major causes of death to the black-white life expectancy gap by US state

**DOI:** 10.1101/140152

**Authors:** Corinne A Riddell, Kathryn T Morrison, Sam Harper, Jay S Kaufman

**Affiliations:** Department of Epidemiology, Biostatistics & Occupational Health, McGill University, 1020 Pine Avenue West, Room 27, Montreal, QC H3A 1A2, Canada.

## Abstract

States have fared differently in their progress towards eliminating the black-white life expectancy gap. Our objective is to describe the pattern of contributions of each of six major causes of deaths to the sex-specific black-white life expectancy gap across states over the last half-century, and identify divergent states.

Using vital statistics and census data, we extracted the number of deaths and population sizes for the years 1969 to 2013, by state, gender, race, 19 age groups, and six major causes of death.

Although mortality from cardiovascular disease has decreased dramatically, its contribution to the life expectancy gap increased over time for men (from 0.9 to 1.2 years), but decreased for women (from 2.4 to 1 years). The contribution of non-communicable diseases to the gap was stable over time for men (approximately 0.4 years) but decreased for women (from 0.7 to 0.2 years), while cancers exhibited an inverted-U trend for men (peaking at 1.1 years in 1988) and a stable contribution for women (approximately 0.5 years). Both genders exhibited a decreased contribution from injuries (men: 2.2 to 0.4 years), that became negative for women (women: 0.5 to -0.1 years). Several states diverged from these general trends.

Life expectancy for both races has improved substantially in the US. For men, much of this improvement was due to narrowing differences in injury-related mortality, but these contributions were rivaled by an increasing gap in CVD-related mortality. In women, a crossover in injury-related mortality led to a narrower gap, realized partially by increasing mortality among whites.

## Significance

Racial differences have been a fixture of American life for centuries, but their continued evolution is a topic of great relevance for public policy and social justice. States have fared differently in eliminating the black-white life expectancy gap. Using vital statistics and census data, we describe how six major causes of death contributed to the life expectancy gap between blacks and whites from 1969 to 2013. We find that states diverged in their cause-of-death specific contributions to the life expectancy gap in ways that either resulted in the near or total elimination of the gap, or alternatively its exacerbation. This analysis provides a foundation for future work to investigate state characteristics that may result in inequalities in mortality by race.

Life expectancy is a barometer for overall population health. In 2014, life expectancy at birth in the United States was 78.8 years, but hidden within this summary are important subgroups differences by self-defined racial ethnicity. Whites lived 3.4 years longer than blacks.^1^ While both black and white Americans have experienced increases in life expectancy since the beginning of the 20 ^th^ century, the difference between their life expectancies has varied.^2^

During the 1980s, the life expectancy gap widened and then narrowed in the 1990s due largely to relative mortality improvements among blacks in deaths due to homicide, HIV/AIDS, unintentional injuries, and, among women, due to differential improvements in heart disease.^3^ The gap continued to narrow during the 2000s, but states differed markedly in the magnitude of improvement. For example, the life expectancy gap in New York decreased from 8.0 to 2.4 years between 1990 and 2009, but in California the decrease was much more modest — from 6.7 to 5.6 years.^4^ However, the extent to which different causes of death may have contributed to state-level changes in the black-white gap are unknown.

The over-arching objective of this study is to describe the contribution of selected causes of death to the black-white life expectancy gap over forty-five years, and identify divergent states and gender differences. To do so, we used publicly available data from 1969 to 2013 detailing the number of deaths and population size according to US state, year, gender, race (black or white), age, and cause of death. We excluded 11 states with the smallest black populations: Alaska, Hawaii, Idaho, Maine, Montana, New Hampshire, North Dakota, South Dakota, Utah, Vermont, and Wyoming. These states contained less than 0.4% of the black population in the United States in 2010.^5^ We did not consider Hispanic ethnicity as part of our racial definition, since Hispanic origin was ascertained beginning in 1978 on death certificates in certain states and 1980 for the Census.^6^

Using these data, we estimated state-level trends in the black-white life expectancy gap measured in years separately for men and women. We then calculated the contribution of each of six major causes of death: cardiovascular disease, cancer, non-communicable disease, injury, communicable disease, and all other causes. This study lays the foundation for the identification of policies or social conditions in highly successful states, and the suggestion of broad targets for interventions to further reduce the gap in less successful states.

## Results

**Figures 1 and 2** display trends in the life expectancy gap for men and women, respectively. For men and women, the life expectancy gap has decreased since 1969, although several states exhibited a period of increasing inequality that peaked in the early 1990s, especially among men.

**Figure 1:**
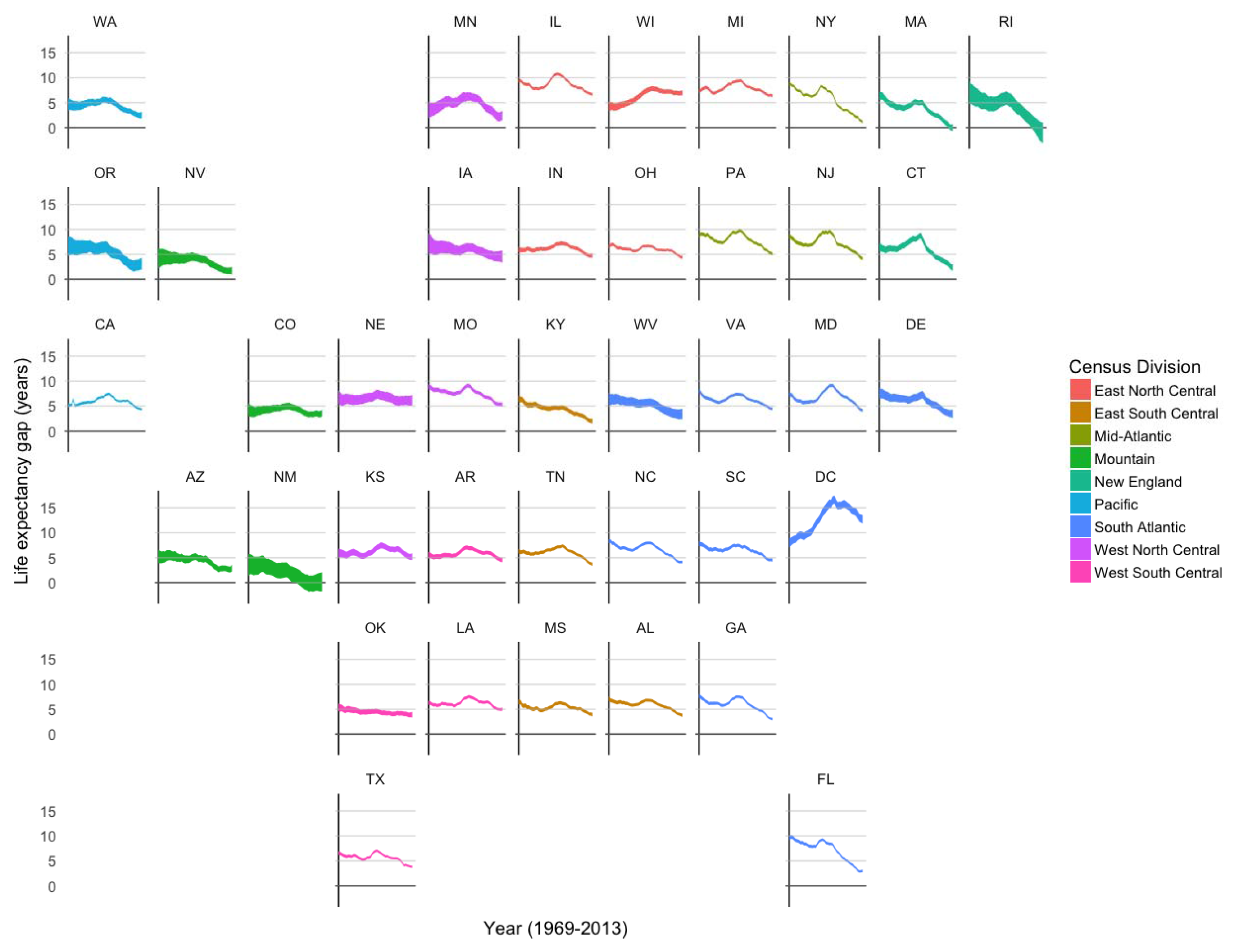
State-level trends in the black-white life expectancy gap in men, United States, 1969–2013

**Figures 2–14** display trends in the contribution of each broad cause of death to the life expectancy gap for men and women, in years, for each state. The blue line depicts the mean of the smoothed contribution estimated for each state, such that states can have very different trend lines. These lines are overlaid with the meta-estimated national trend, shown with a red line. We supplement these main results with smoothed estimates of the age-standardized mortality rates for each cause of death (**Figures S1-S12**).

**Figure 2:**
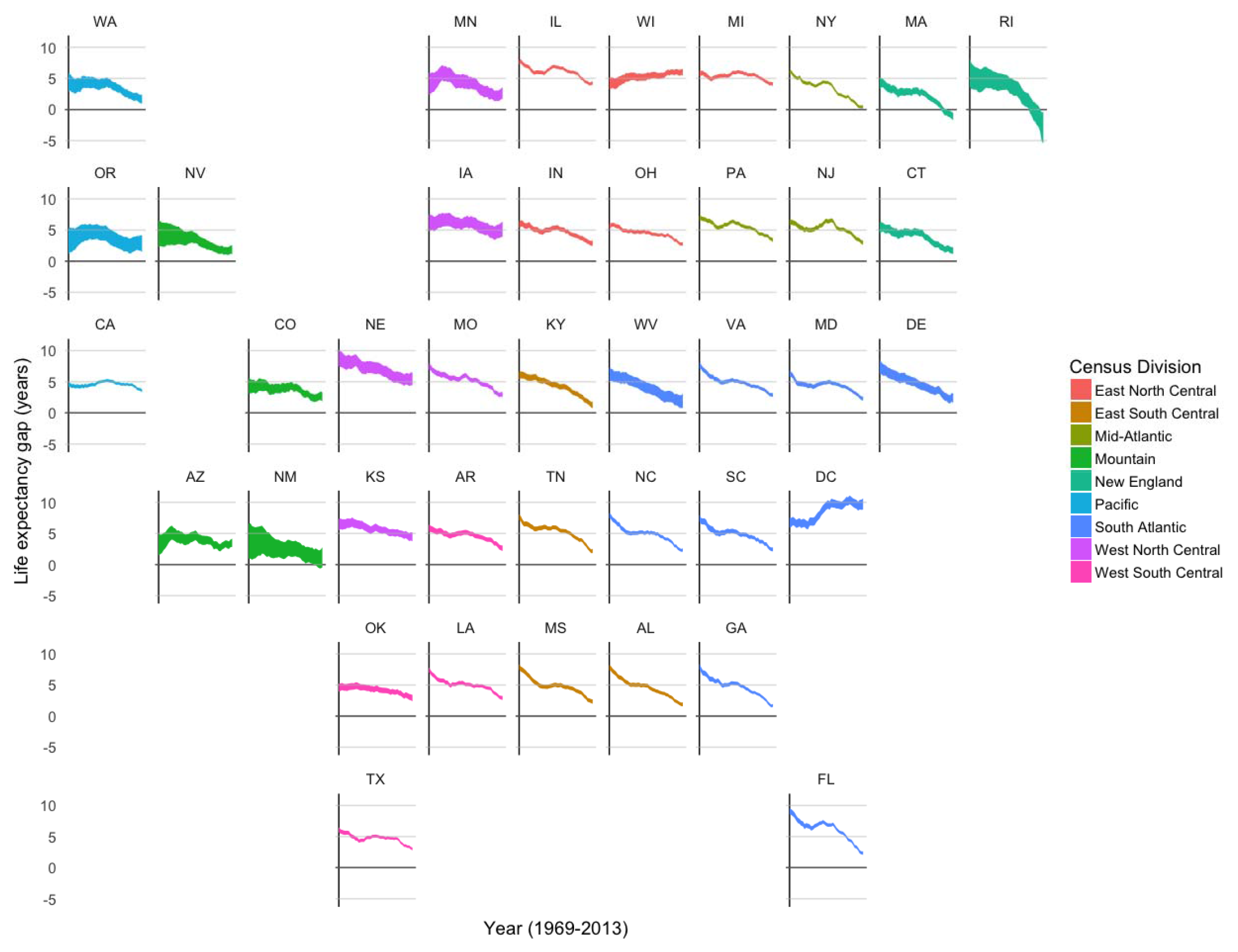
State-level trends in the black-white life expectancy gap in women, United States, 1969–2013

### Cardiovascular disease

Cardiovascular disease (CVD) is the most common cause of death for both men and women of both races. Mortality from CVD decreased markedly since 1969 for all groups (**Figures S1 and S2**), but white men underwent relatively larger mortality decreases than black men. As a result, the contribution of CVD to the black-white life expectancy gap among men increased modestly for most states (**Figure 3**). Black women experienced relatively larger mortality decreases over time compared to white women, resulting in a decrease to the CVD contribution to the life expectancy gap that was greater than one year in several states (**Figure 4**). For both genders, Michigan, Wisconsin, California, and DC demonstrated higher-than-average increases in the contribution of CVD over time, whereas Massachusetts and New York showed persistently lower CVD contributions.

**Figure 3:**
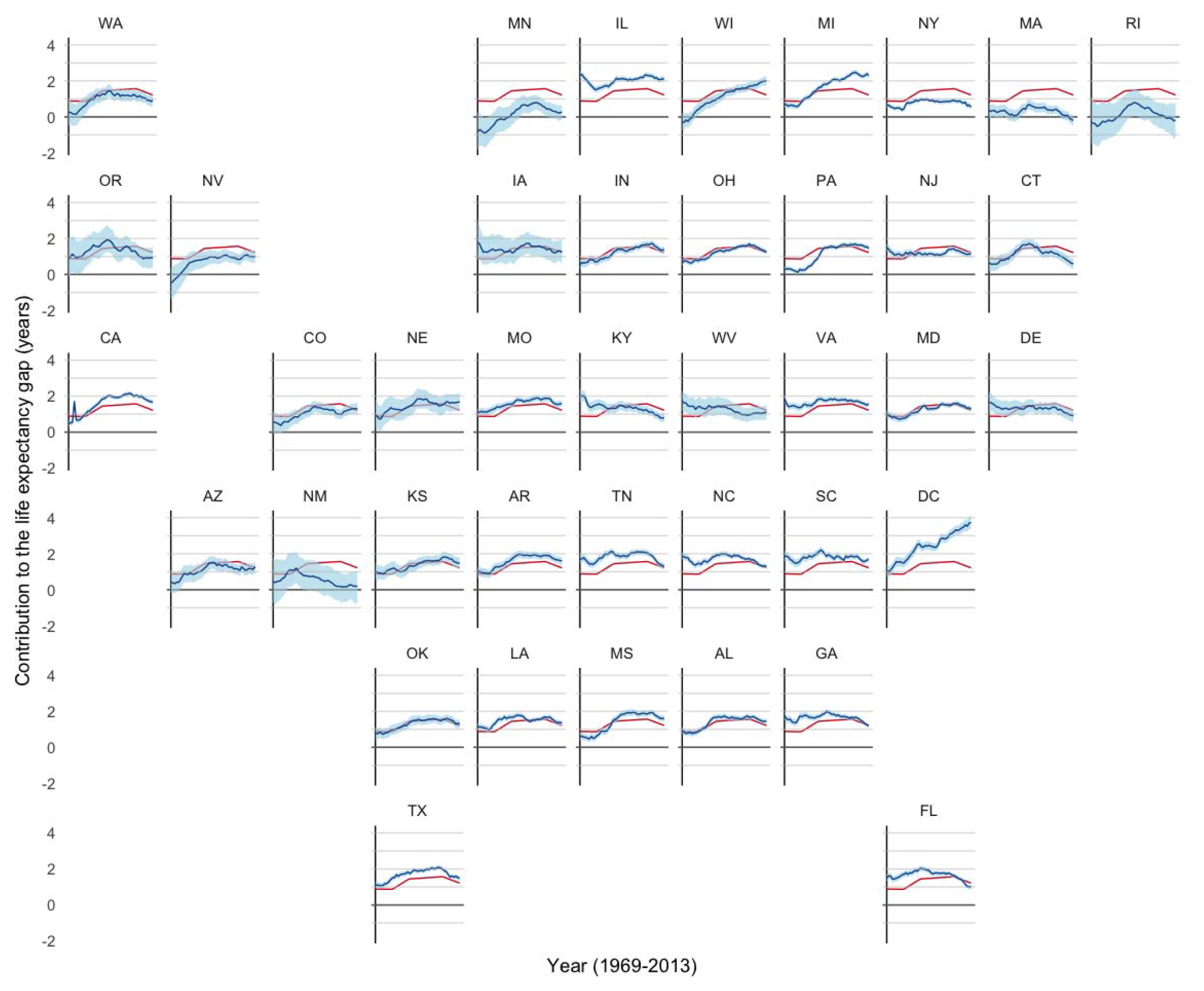
Smoothed state-level trends in the contribution of CVD to the life expectancy gap vs. the national pattern in men, United States, 1969–2013

**Figure 4:**
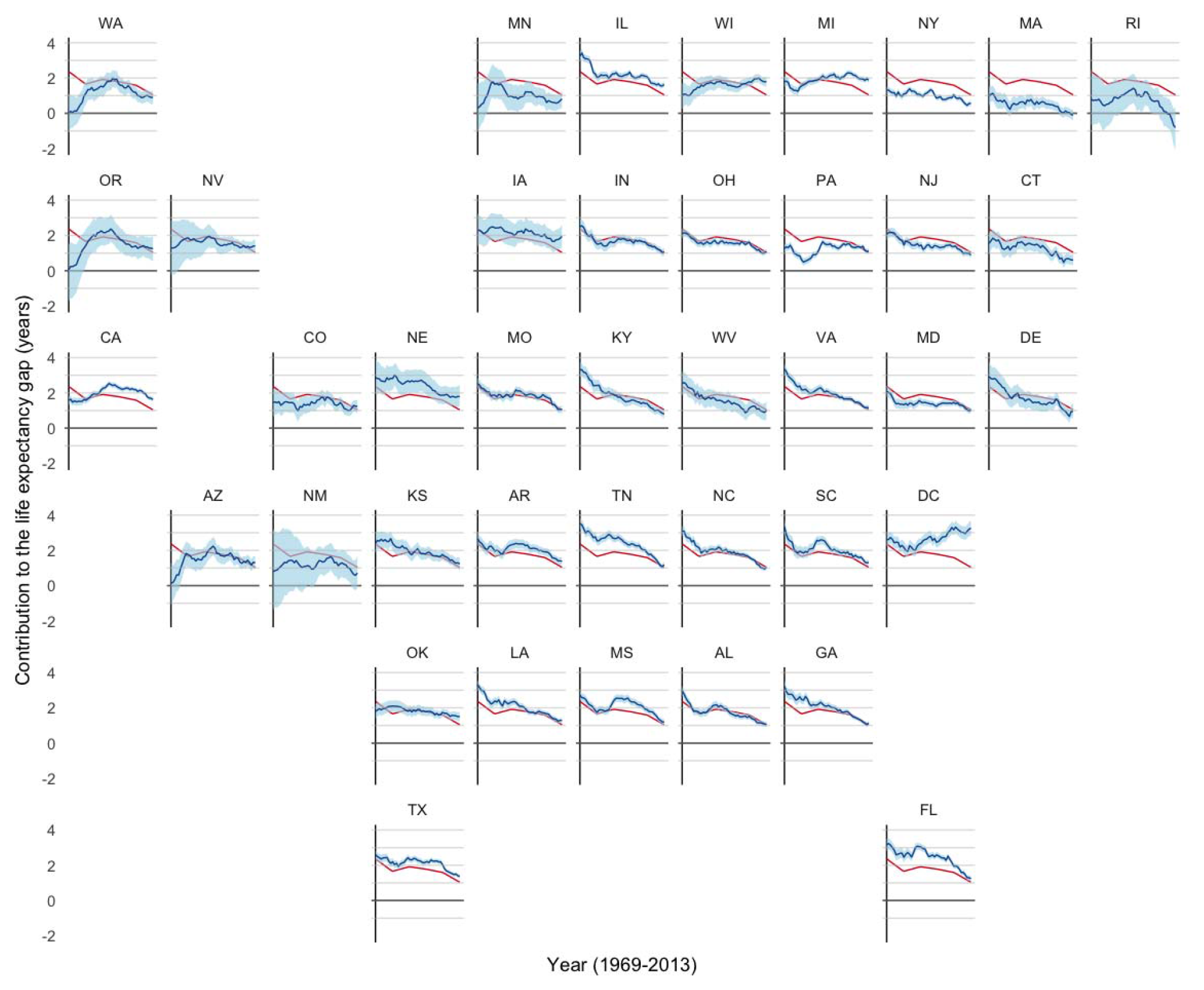
Smoothed state-level trends in the contribution of CVD to the life expectancy gap vs. the national pattern in women, United States, 1969–2013

### Cancer

For all groups, mortality from cancer peaked in the mid 1990s and decreased thereafter (**Figures S3 and S4**). Among men, this led to an inverted-U shape pattern in cancer’s contribution to the life expectancy gap (**Figure 5**). For women, although cancer-related mortality continues to be higher among blacks in many states, the mortality trends were similar between blacks and whites, resulting in a flat general pattern in the contribution of cancer to the life expectancy over time (**Figure 6**). For both genders, DC and Wisconsin exhibit the largest cancer contributions (and also exhibit high initial values among men), while New York and Massachusetts experienced large decreases in the contribution of cancer to the life expectancy gap.

**Figure 5:**
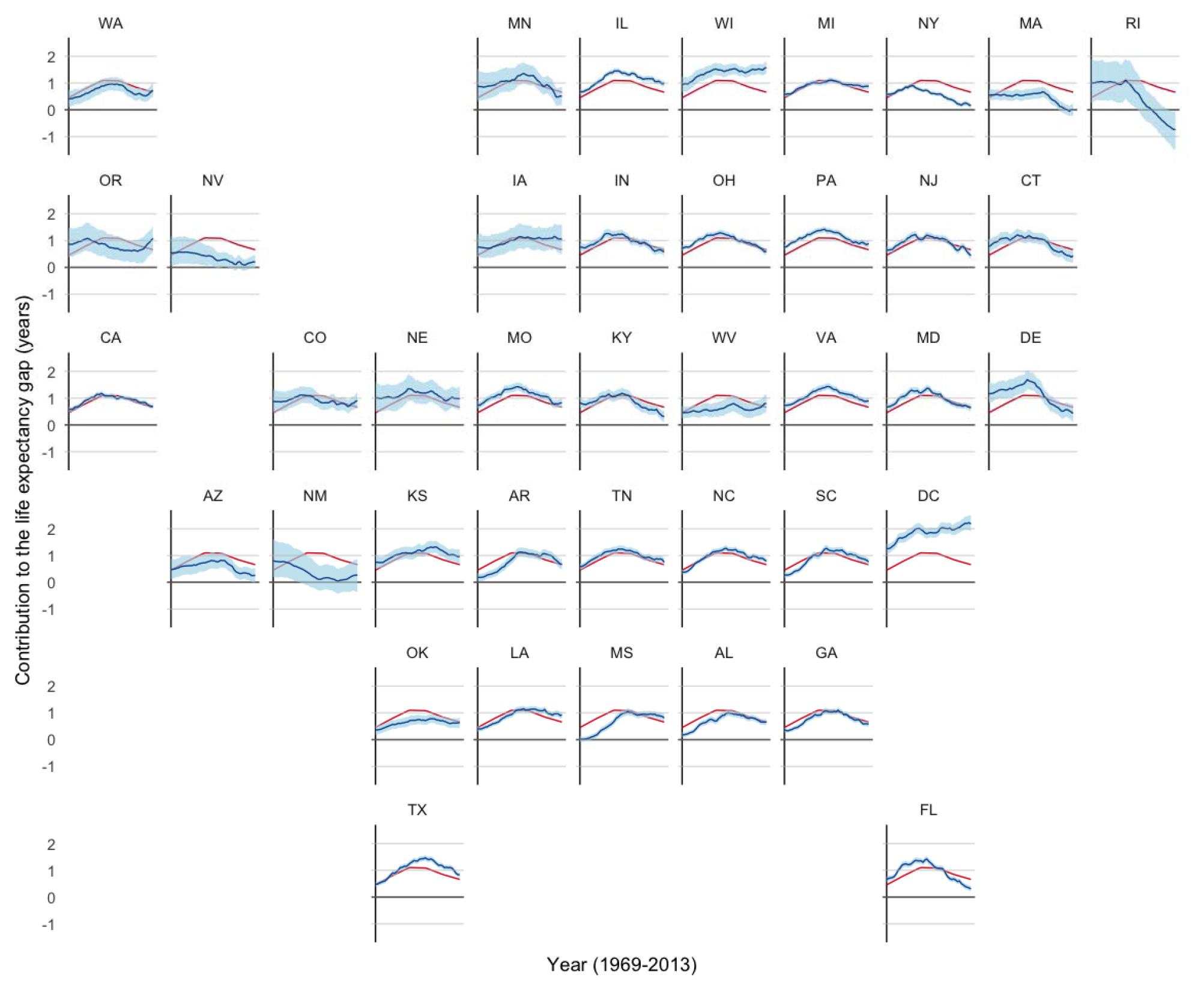
Smoothed state-level trends in the contribution of cancer disease to the life expectancy gap vs. the national pattern in men, United States, 1969–2013

**Figure 6:**
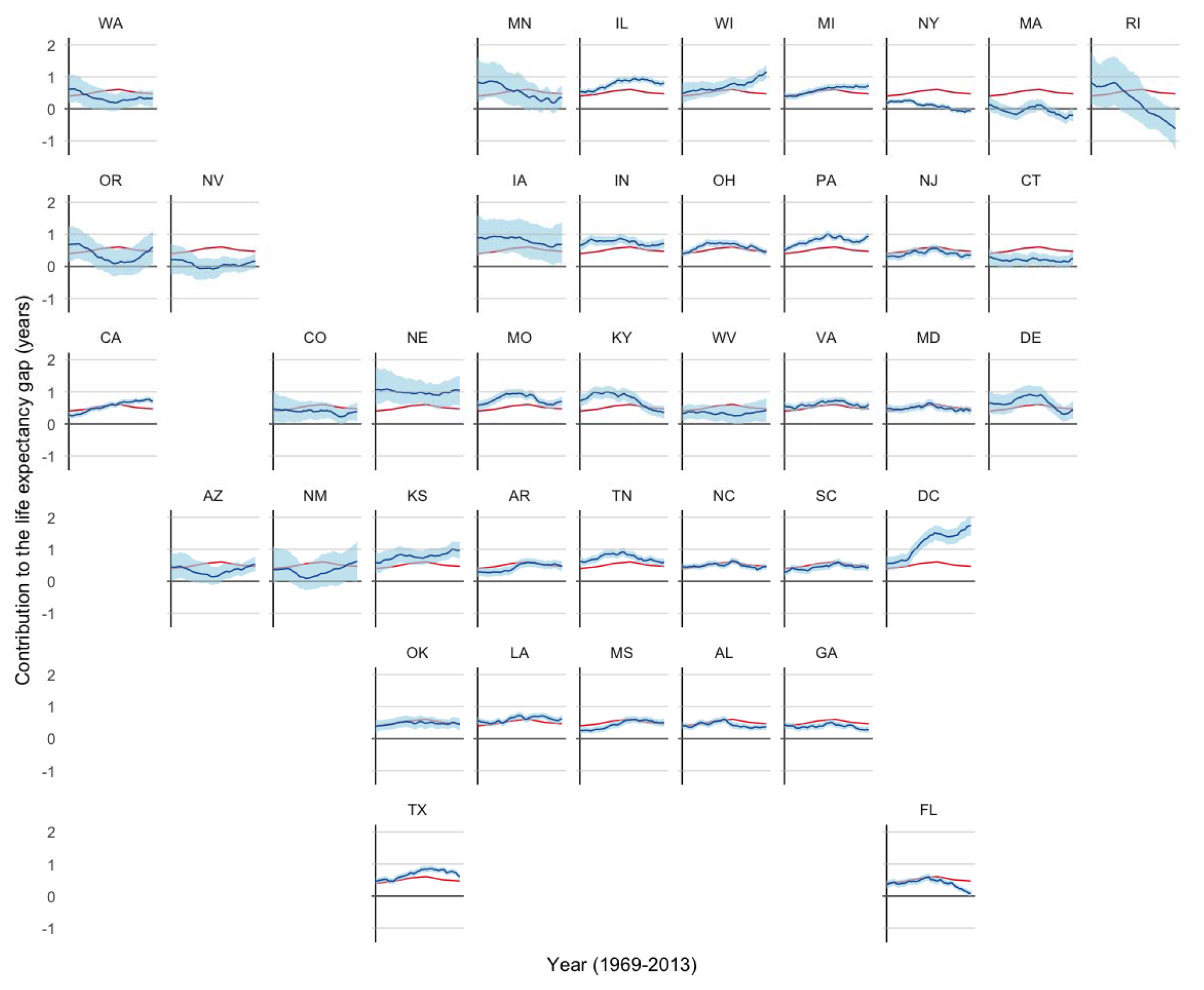
Smoothed state-level trends in the contribution of cancer disease to the life expectancy gap vs. the national pattern in women, United States, 1969–2013

### Non-communicable disease

Mortality from non-communicable disease increased over time for men in most states, with DC as a notable exception **(Figure S5)**. Recently, there was a slight decrease in mortality rates, especially among black men. These trends are similar among women, except that black women exhibited a sharper decline in mortality in recent years, eliminating the mortality difference in several states **(Figure S6)**. As such, the contribution of non-communicable disease to the life expectancy gap was generally flat among men with a recent decrease, and exhibited a gradual decrease over time among women **(Figures 7 and 8)**. As with other causes, New York and Massachusetts exhibited decreasing contributions, and Nevada also made substantial progress. Furthermore, while DC’s mortality rates decreased, they presently have the highest contribution of non-communicable disease to the gap.

**Figure 7:**
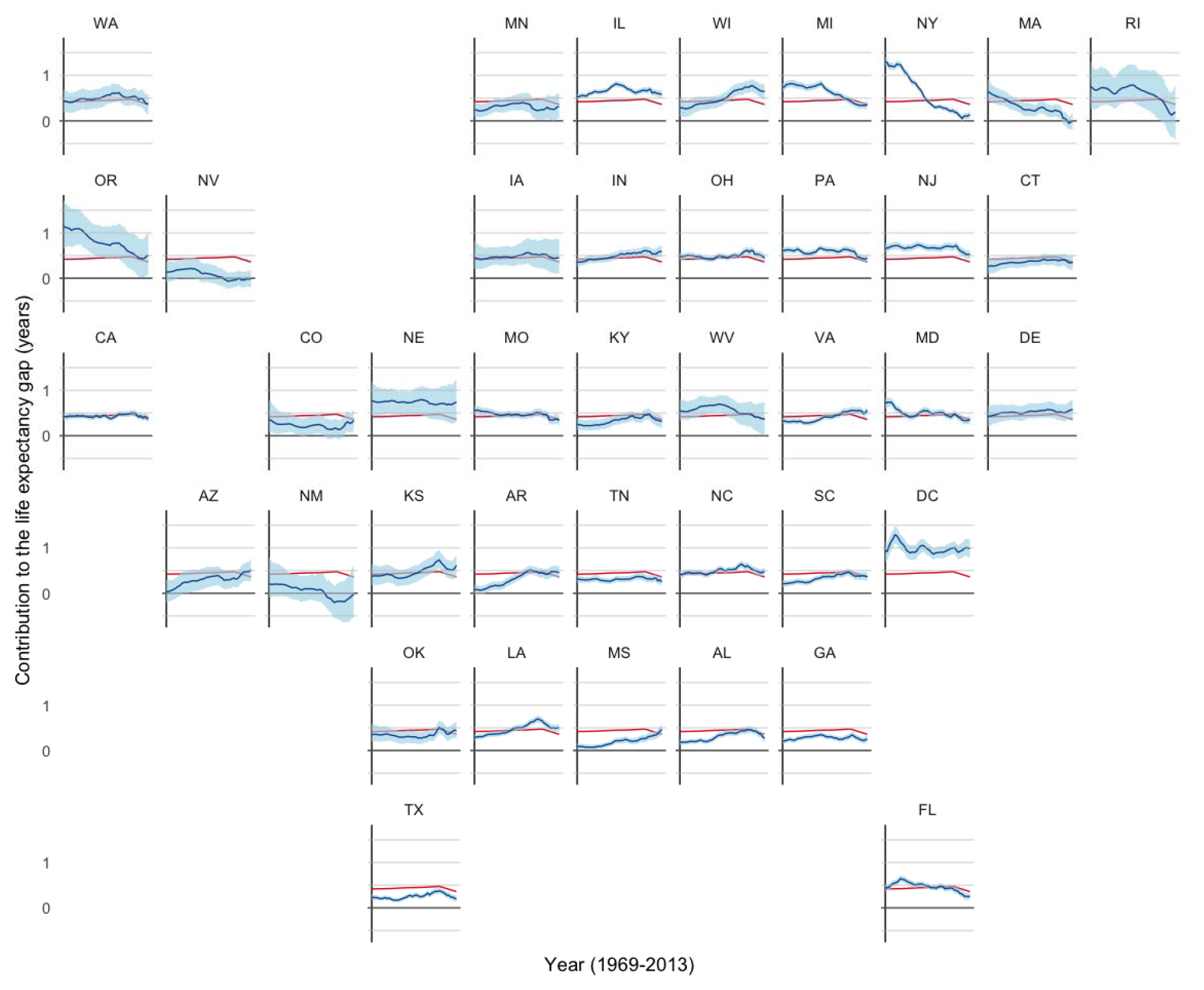
Smoothed state-level trends in the contribution of non-communicable disease to the life expectancy gap vs. the national pattern in men, United States, 1969–2013

**Figure 8:**
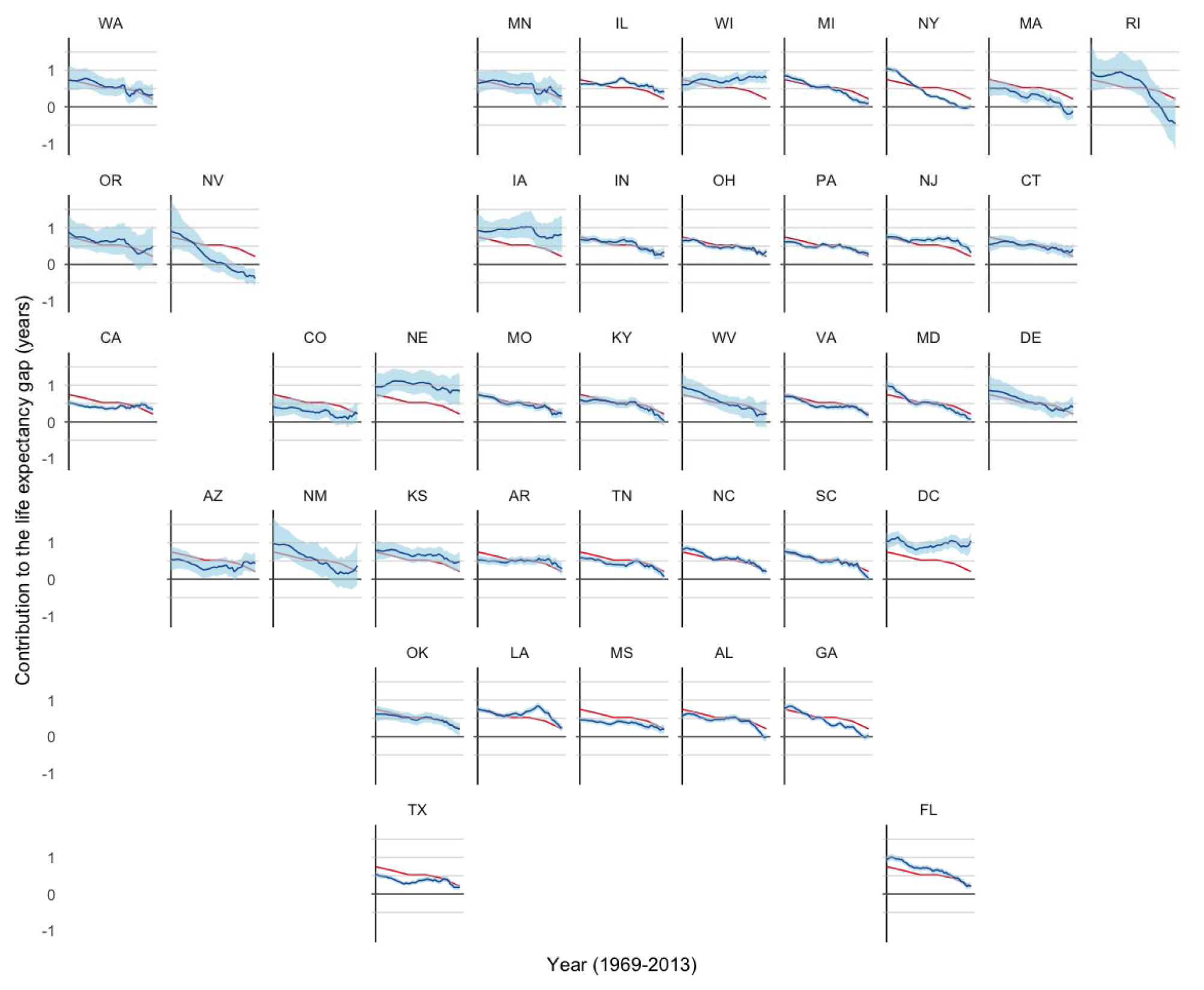
Smoothed state-level trends in the contribution of non-communicable disease to the life expectancy gap vs. the national pattern in women, United States, 1969–2013

### Injury

Trends in mortalilty from injuries differ for blacks and whites. Over time, injury-related mortality has decreased precipitously among blacks, although an epidemic of homicide resulted in heightened mortality from injuries among black men during the 1980s and early 1990s **(Figure S7).^3^** For whites, most states exhibit an initial period of gradual decreases in injury mortality, followed by a period of gradual increases, especially among white women **(Figure S8)**. Thus, injury mortality rates among blacks are equivalent, or lower than those among whites in many states, with white women demonstrating sharper recent increases. The contribution of injuries to the mortality gap has therefore generally decreased over time, and has become negative in those states that now have lower injury-related mortality among blacks than whites **(Figures 9 and 10)**. Among men, DC, Illinois, Michigan and Missouri exhibit the largest recent injury-related contributions. Among women, many southern states exhibit negative contributions (indicating lower injury-related mortality among black women), while DC, Illinois, and Michigan having higher contributions than the general trend.

**Figure 9:**
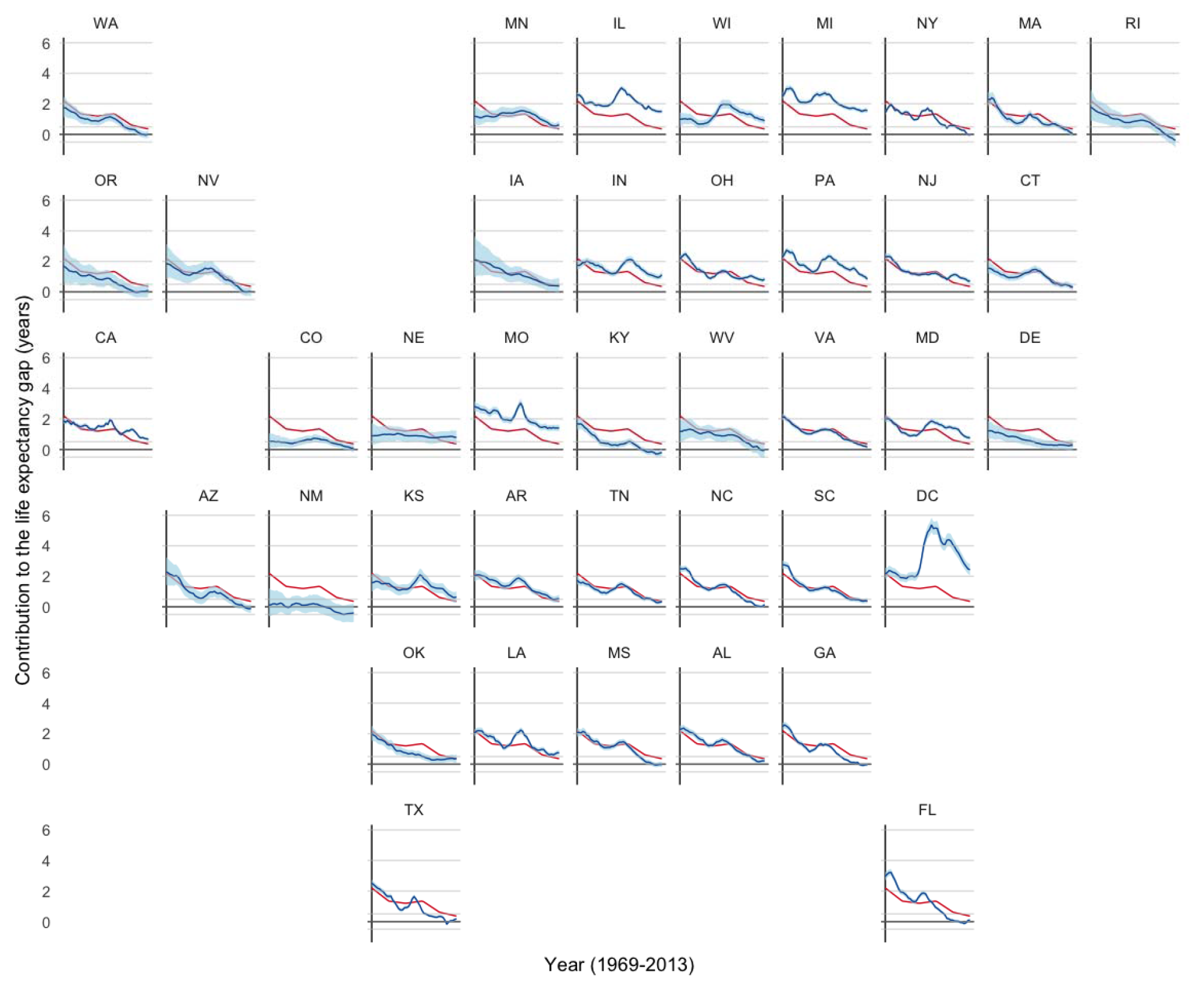
Smoothed state-level trends in the contribution of injuries to the life expectancy gap vs. the national pattern in men, United States, 1969–2013

**Figure 10:**
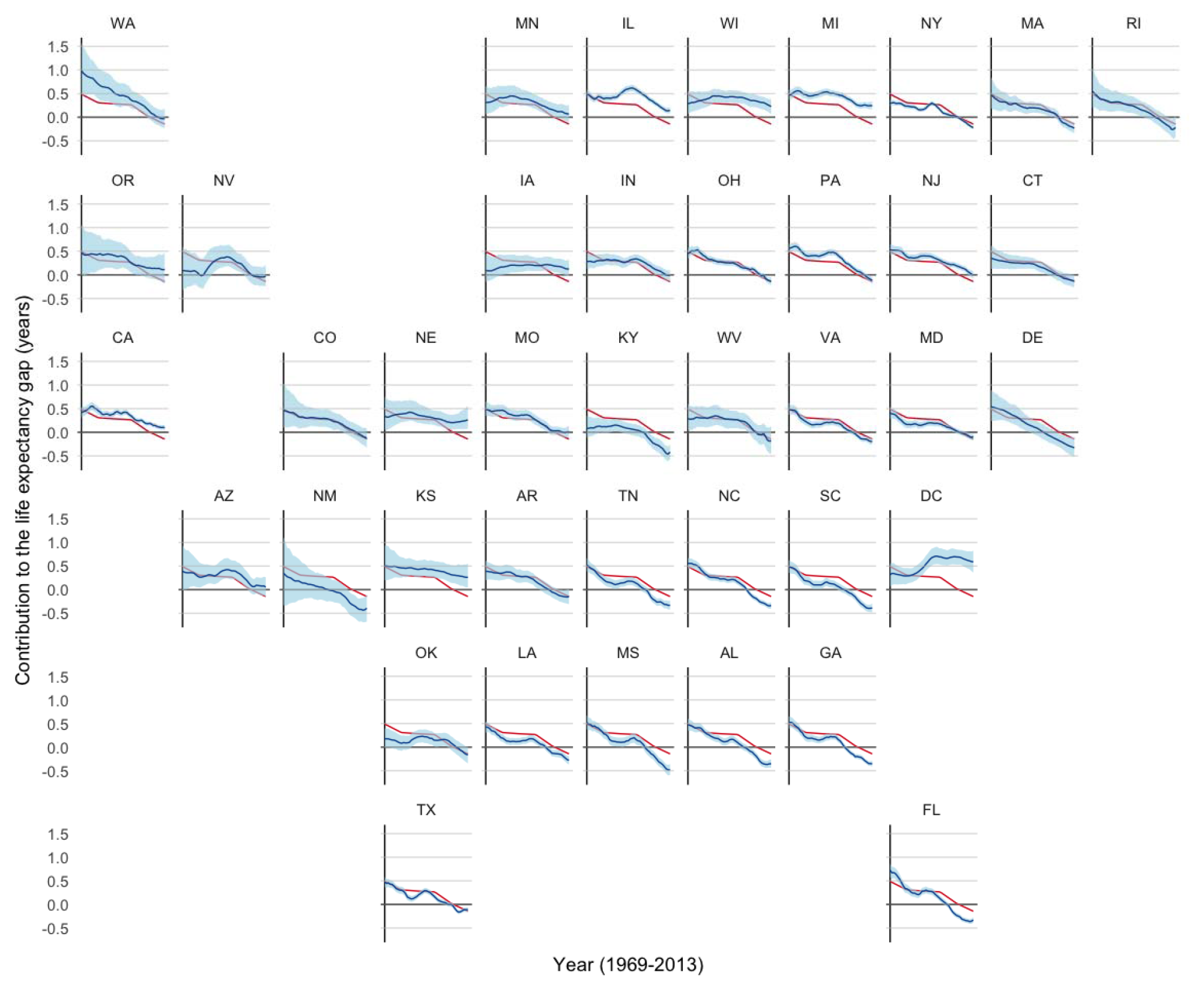
Smoothed state-level trends in the contribution of injuries to the life expectancy gap vs. the national pattern in women, United States, 1969–2013

### Communicable disease

Of the major causes that we considered, mortality from communicable disease was the lowest, although the 1990s exhibited an enormous increase in mortality at the height of the HIV/AIDS epidemic before antiretroviral therapy was available **(Figures S9 and S10)**. This spike was greater among men, especially blacks, and among states with larger urban centers. The contribution of communicable disease to the life expectancy gap therefore exhibited a peak in the mid 1990s, with states diverging from this general pattern according to the intensity of their local epidemics **(Figures 11 and 12)**. DC still exhibits higher communicable disease-related contributions among men, while New Jersey and Florida additionally show higher contributions among women.

**Figure 11:**
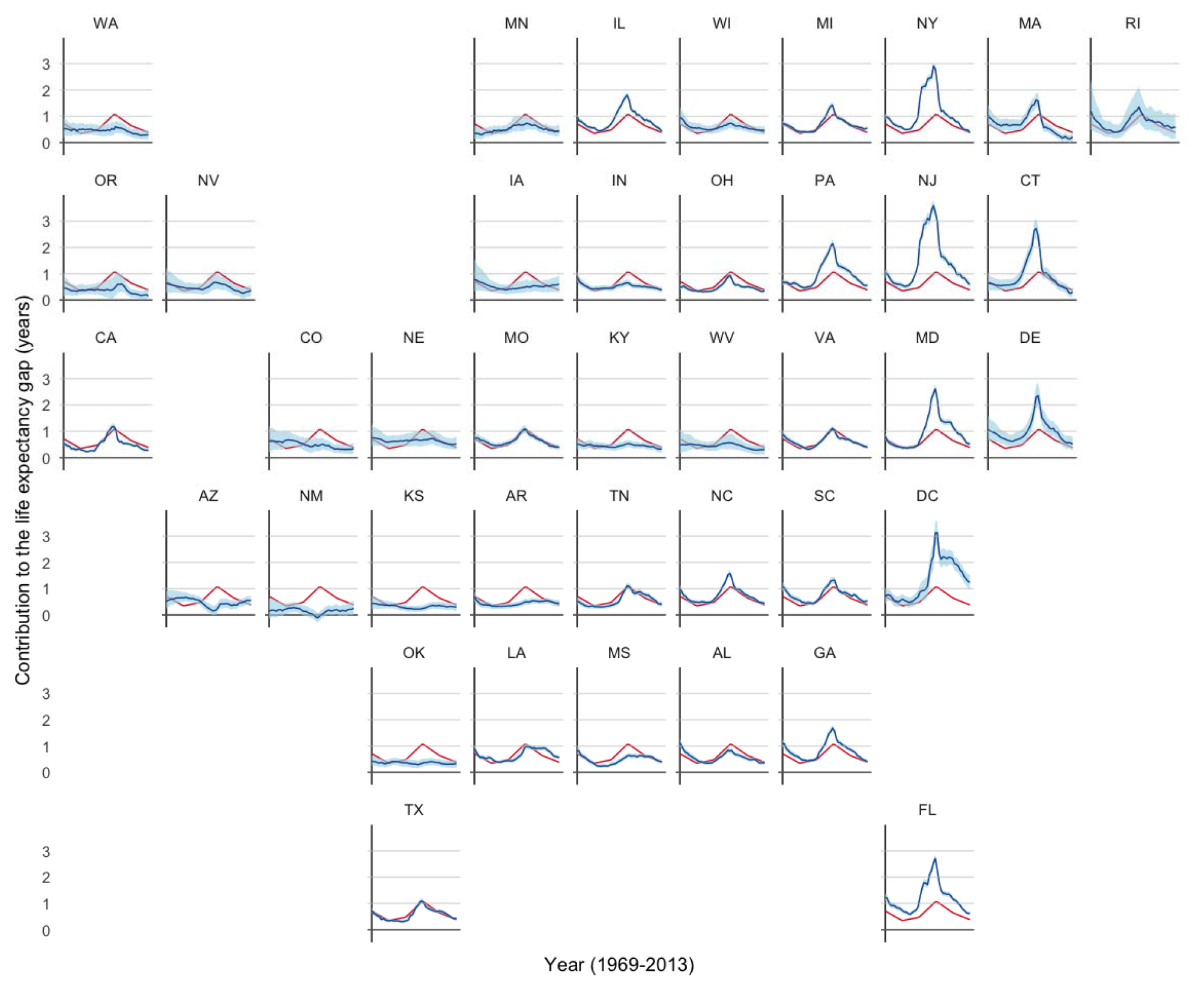
Smoothed state-level trends in the contribution of communicable disease to the life expectancy gap vs. the national pattern in men, United States, 1969–2013

**Figure 12:**
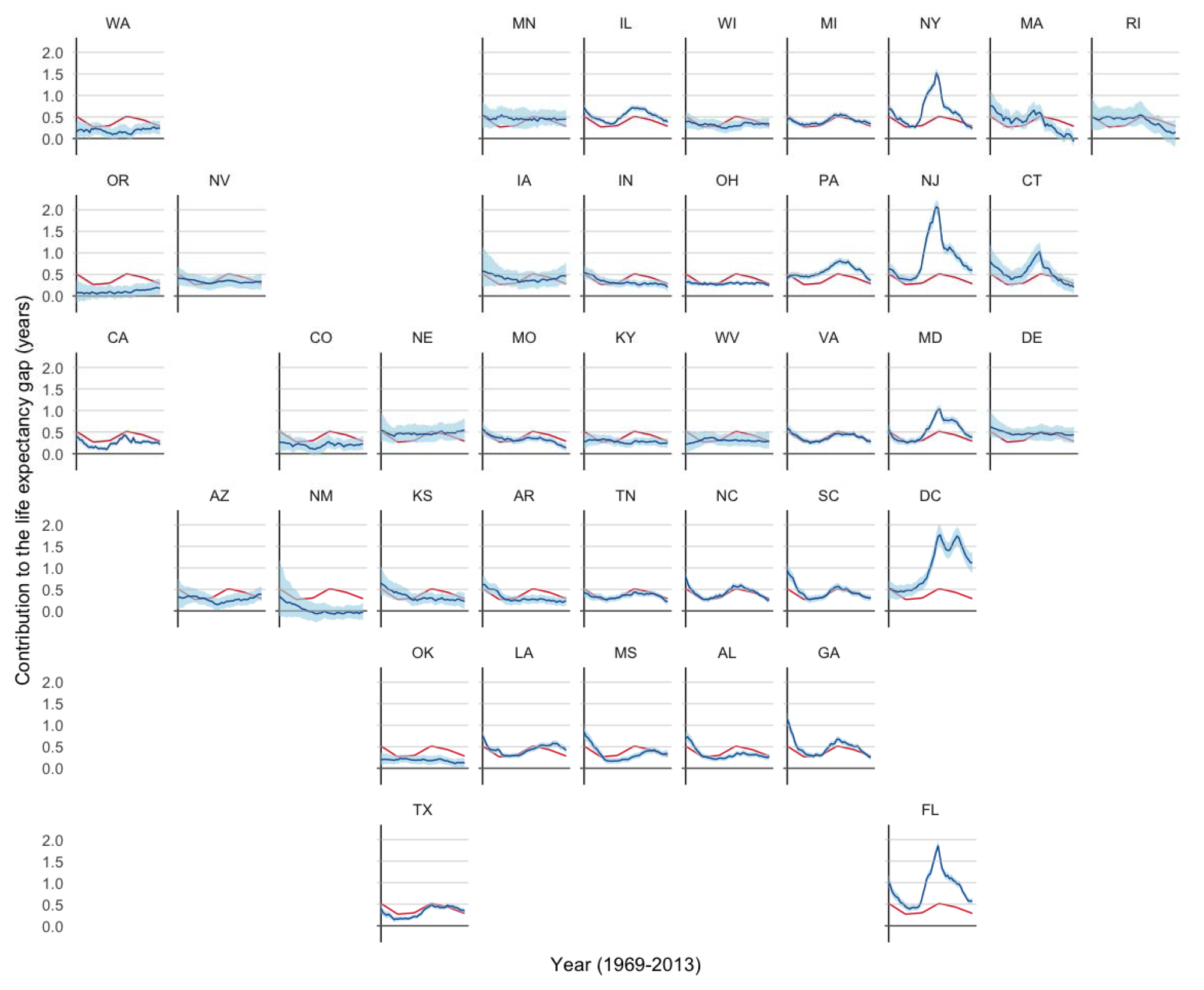
Smoothed state-level trends in the contribution of communicable disease to the life expectancy gap vs. the national pattern in women, United States, 1969–2013

### All other causes

Mortality from all other causes generally decreased earlier in the time period, especially among blacks (**Figures S11 and S12**). White men and women have shown increasing mortality rates from all other causes since the 1980s, along with black women, whereas black men have continued to experience decreasing rates within several states. As such, the contribution of all other causes to the life expectancy gap has generally decreased over time, and is virtually zero in New York, Massachusetts, and Rhode Island for both genders (**Figures 13 and 14**). The most disparate states are Pennsylvania and DC, with much larger contributions estimated for all other causes in the earlier years within Pennsylvania and in the later years within DC.

**Figure 13:**
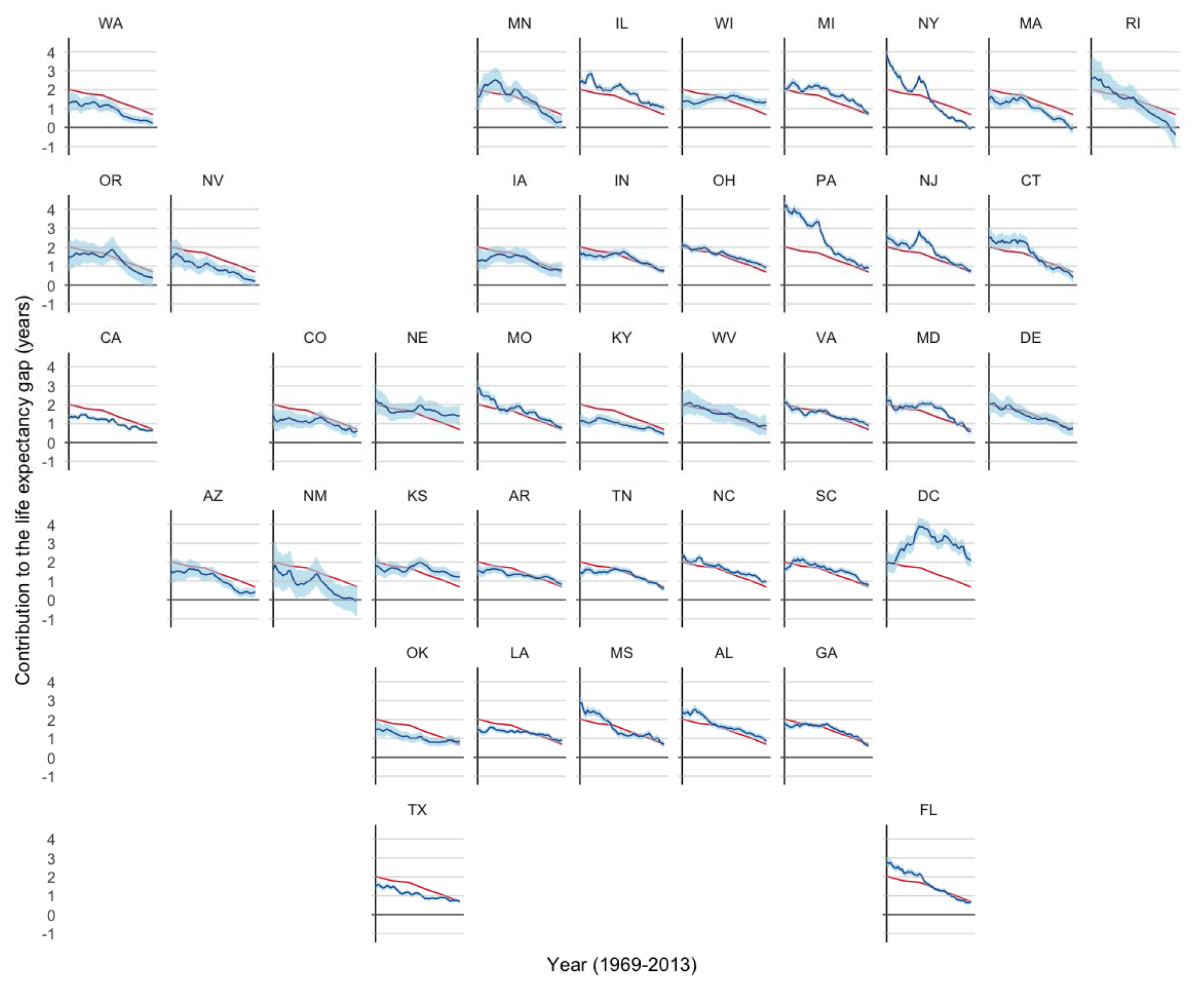
Smoothed state-level trends in the contribution of all other causes to the life expectancy gap vs. the national pattern in men, United States, 1969–2013

**Figure 14:**
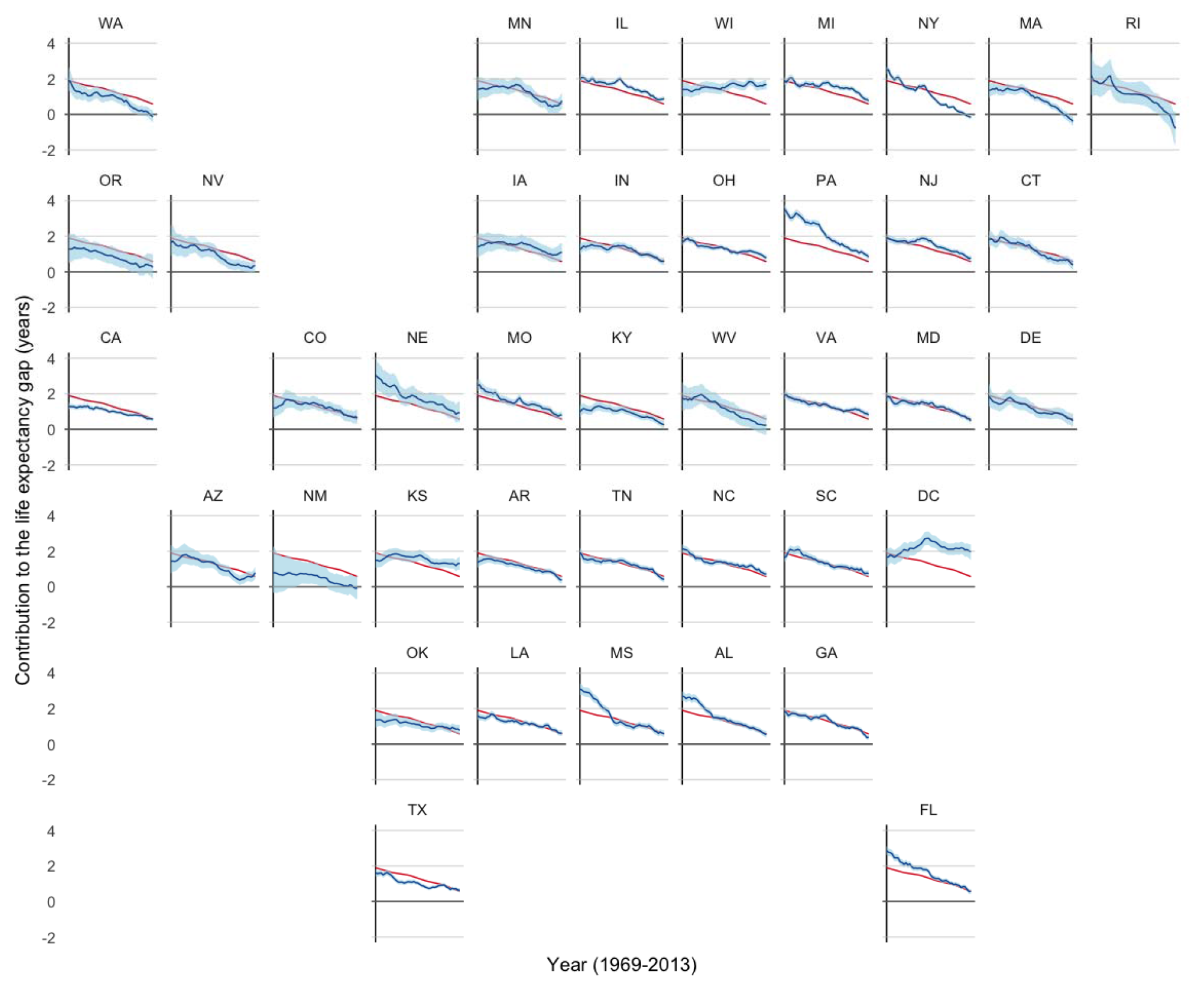
Smoothed state-level trends in the contribution of all other causes to the life expectancy gap vs. the national pattern in women, United States, 1969–2013

Because the majority of infant deaths are classified as being due to all other causes in our analysis, we also tried to separate any impact of all other causes from those of infant mortality more generally. We conducted a sensitivity analysis to calculate the contribution of deaths due to all other causes that occurred during the first year of life to the life expectancy gap (Figures 15 and 16). We found that 43% of the contribution of all other causes to the gap was due to differences in infant mortality in 1969. In 2013, this proportion reached 53%. Thus, the trend in the all other causes contribution partially reflects a decreasing contribution from infant mortality.

**Figure 15:**
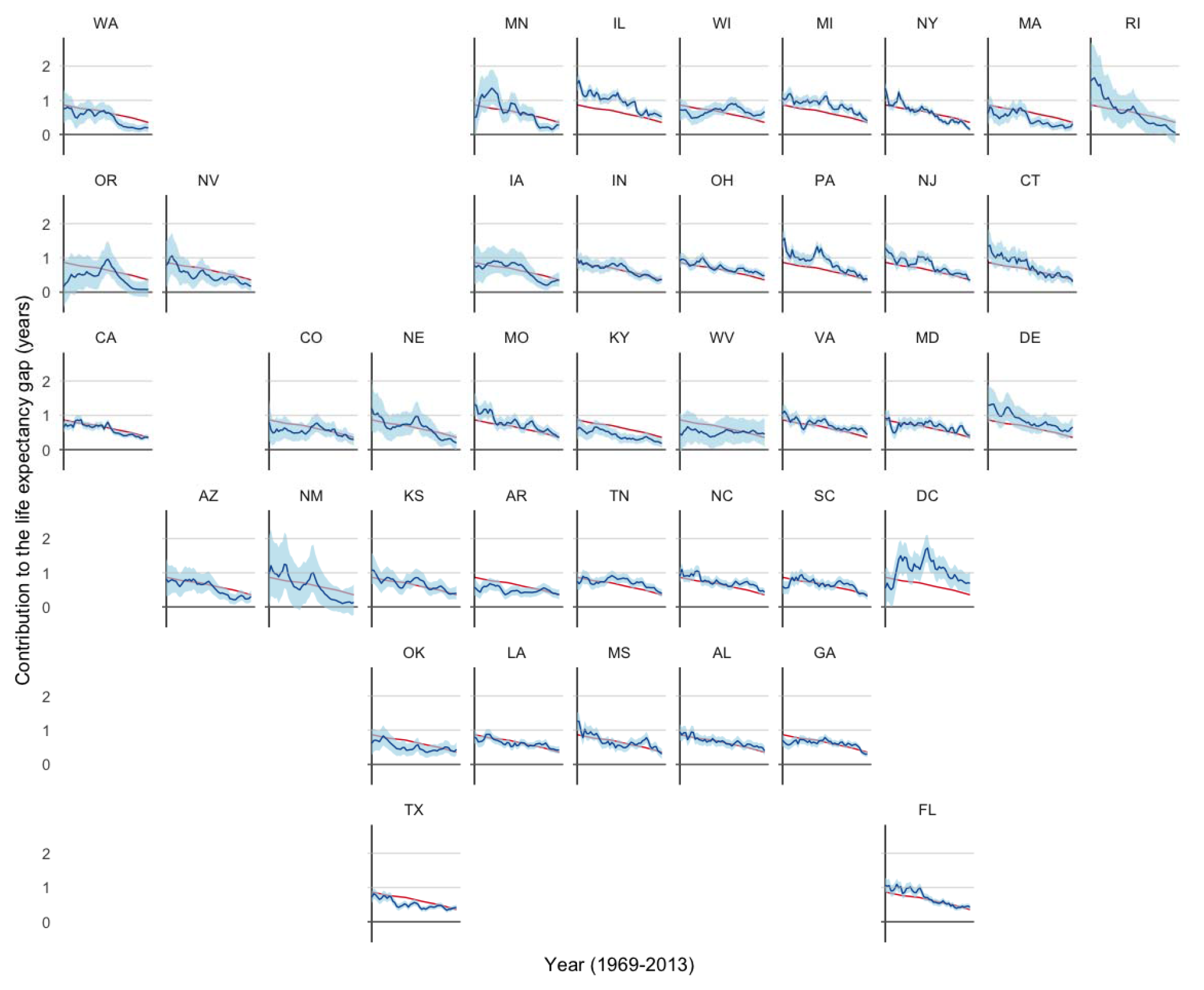
Smoothed state-level trends in the contribution of all other causes to the life expectancy gap vs. the national pattern in men younger than one, 1969–2013

**Figure 16:**
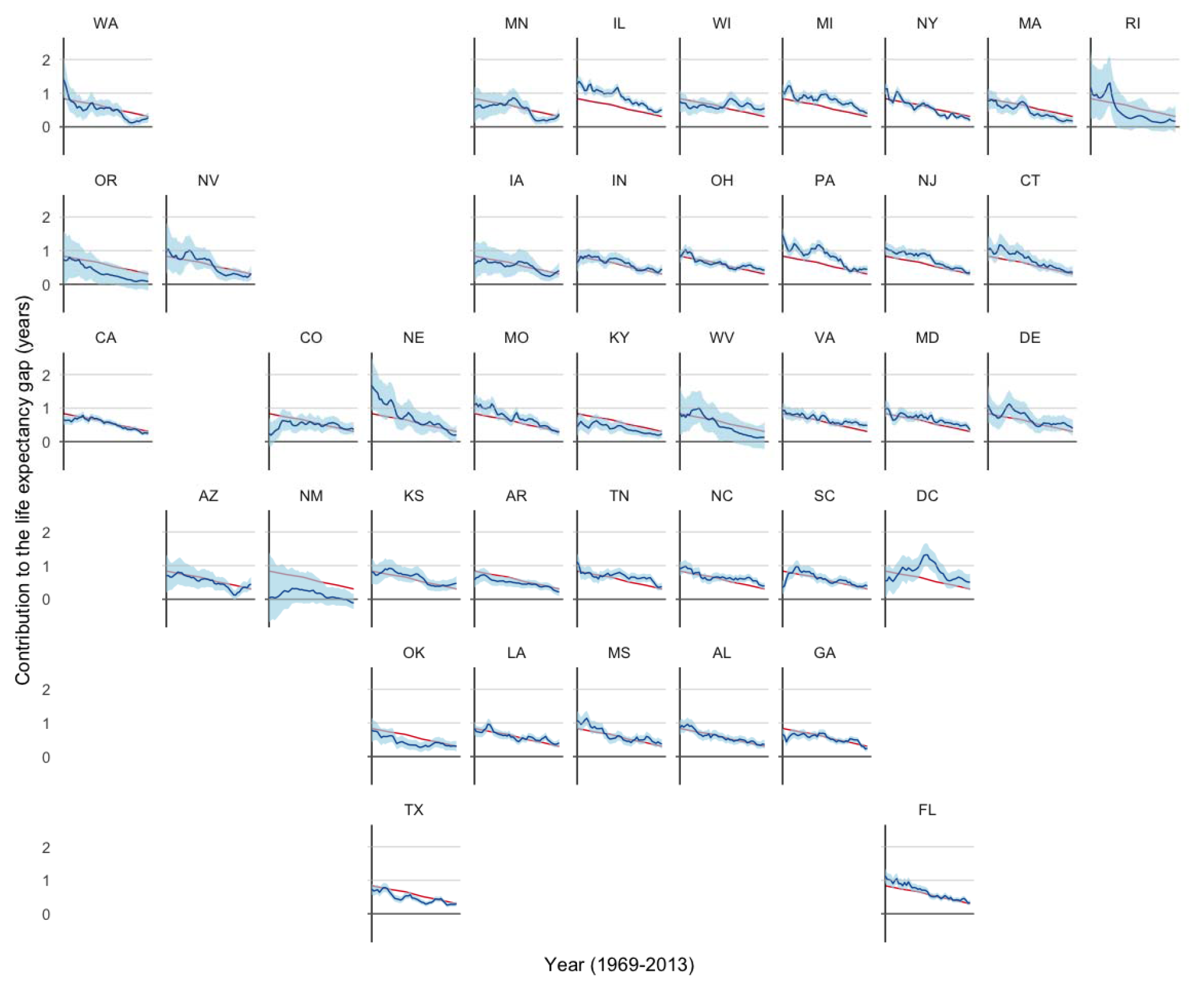
Smoothed state-level trends in the contribution of all other causes to the life expectancy gap vs. the national pattern in women younger than one, 1969–2013

## Discussion

### Summary of findings

We examined state-level trends in six major cause-of-death contributions to the black-white life expectancy gap over a 45 year period. While we found that many states conformed to the national pattern of cause-of-death specific contributions, several states diverged in ways that either resulted in the near or total elimination of the life expectancy gap, or alternatively its exacerbation. In contrast to some recent work focused on county-level inequalities,^7,8^ states have some substantial advantages as analysis units; They have large populations to generate stable estimates, tend to be more demographically stable over time, and they are policymaking entities, and so have divergent social agendas that are reflected in varying patterns of mortality by race.

Overall, the racial gap in life expectancy is now smaller in many states in the South and Southeast that are associated historically with greater degrees of inequality and slower social progress, compared to the mid-West, which had been the receiving point of “the Great Migration” for improved economic opportunities in the mid-20^th^ Century.^9^ Trends in the black-white life expectancy gap among women in “rust-belt” states such as Michigan, Wisconsin and Illinois appear especially discouraging, compared to remarkable progress in Florida, North Carolina, Georgia and Mississippi. On the one hand, it appears to run counter to the narrative of worse overall health progress in the former Confederacy.^8^ On the other hand, greater racial equality can be achieved by falling black mortality rates or rising white rates, and so the closing gap cannot be celebrated without a careful examination of the two individual trends.^10^

Consistent with what has been reported at the national level,^11^ we found that CVD makes the largest contribution to the life expectancy gap, even though both races and genders experienced decreased CVD mortality rates over time. Injuries (especially homicide) continue to play a role in contributing to the current gap for men in several states, especially in the Midwest and DC, while contributing to a decreased gap among women in the South. This decreased gap among women likely reflects the higher rate of mortality from drug overdoses among whites compared to blacks during the ongoing opioid epidemic.^12^ Communicable disease continues to contribute to the gap in most states, especially in DC, New Jersey, and Florida. In our previous work, approximately half of the contribution of communicable disease to the gap was found to be due to HIV in 2008 nationally,^11^ suggesting that these states may increase equality through interventions targeted at reducing mortality from HIV. Cancer is also responsible for large contributions to the life expectancy gap in some states (Illinnois, Michigan, and Wisconsin) but not in others (New York, Massachusetts). Overall, New York and Massachusetts have virtually eliminated the black-white life expectancy gap for both men and women.However, both states began with only a minimal contribution of CVD and cancer to the mortality gap. The difference in early contributions is especially stark for the contribution of CVD among women in 1969, where many states, especially those in the South, had CVD contributions of more than two years. Thus, the reason for New York’s and Massachusetts’ low CVD contributions does not appear to be the effect of recent changes, but highlights a long-standing pattern of greater equality and points to the importance of viewing the most recent estimates in light of historical patterns.

The most important implications for a state-level analysis are using the patterns to make connections between changes to social policy and health outcome changes over time. States pursue varying approaches to an array of relevant programs and laws including tobacco control,^13^ Medicare coverage,^14^ hand-gun violence,^15^ and nutrition.^16^ These presumably affect health and longevity, and can have especially important impacts on racial differences when the policies are targeted to disadvantaged groups or outcomes that differentially affect blacks.^17^ The analyses reported here are entirely descriptive, but should be used for subsequent work that seeks to identify the causal processes that link social and health policies to successful trajectories in some states, and worsening disparities in others. Variation in policies over time can be used in quasi-experimental designs to uncover evidence for the role of specific policy innovations, which is an advantage of using states as the units of analysis as opposed to larger regions or smaller counties, neither of which generate policy variations.^18^

#### Strengths and weaknesses

We used data over 45 years, across two racial groups, and for all ages. By examining a wide time window, trends were examined that provided context for the contributions estimated in the most recent year. This allowed us to recognize when the estimated contributions for a particular cause of death were relatively low because they started low at the beginning of the follow-up, versus being low because they had decreased over time. Furthermore, our analysis was centered on decomposing life expectancy at birth, a measure that integrates mortality rates across all ages. This is especially important in light of recent studies that have received criticism for narrowly focusing on a specific age group.^19,20^ The Arriaga decomposition method accounts for the fact that deaths occurring in one cause stratum are reflected in the absence of that death in another stratum. Alongside the decomposition results and age-standardized mortality rates, the scope of our analysis allows readers to examine differences in state-level inequality from many angles.

Our analysis has three main limitations. Firstly, we did not account for Hispanic ethnicity, implying that both black and white populations in our study include some portion of Hispanics that is known to differ over time and by state. This may be especially consequential for blacks in states like Florida or New York that received many black Hispanic immigrants from Cuba, Puerto Rico and the Dominican Republic, potentially changing the composition of the state’s black population to some extent during the study period.^21^ But it also has implications for the white population, especially in states with demographic shifts due to immigration from Mexico such as Florida, California, and Texas. Given that Hispanics have higher life expectanies than non-Hispanics, including Hispanics in the life expectancy calculations will lead to estimates that are likely higher than they would have been had Hispanics been excluded.^22^ We did not to adjust for Hispanic ethnicity because reporting began only in 1978 and even then, agreement between self-reported ethnicity on the Census and secondary-reported ethinicity on the death certificate has been shown to be poorer during the earlier time period and differential according to geography, which could induce measurement error into mortality and disparity estimates.^6^ In a previous sensitivity analysis, we found that accounting for Hispanic ethnicity did not have a substantial effect on the results,^4^ providing some evidence that not accounting for Hispanic ethnicity should not substantially affect our findings. As the US population becomes increasingly Hispanic over time, however, this distinction grows more salient.^23^

Secondly, our analysis does not account for immigration by the foreign-born, or migration of native-born individuals between states. Individuals who immigrate to the US have been shown to have life expectancies at birth that are 5.8 and 4.8 years longer for men and women, respectively in 2008–2010.^24^ As we are examining the difference in life expectancy between blacks and whites, this gap will be affected by the amount of immigration to the extent that new immigrants identify as black or white. Migration between states can also affect the estimated life expectancy of a state if it is associated with underlying risks of death. Migration is particularly important for the District of Columbia, because of it’s small geographical bounds, and substantial in-migration from New York and in- and out-migration to Maryland and Virginia. Middle and upper-class black residents have been shown to move to the suburbs in Maryland and Virginia, while affluent whites are migrating to DC. Both changes would serve to increase the racial gap in DC.^25^

Thirdly, by using broad causes of death we are unable to investigate trends in more specific causes. For example, injury mortality includes homicide, suicide, and drug overdoses, which are known to vary by sex, race, and over time. Drug overdose as a more specific cause of death has played an increasingly important role in shaping disparities in many states, but this is not separately identifiable in our analyses.^26,27^

## Conclusions

Racial differences have been a fixture of American life for centuries, but their continued evolution is a topic of great relevance for public policy and social justice. The careful surveillance of these patterns is therefore a crucial public health activity, and this paper presents a unique perspective on state-level variation over nearly half a century. This analysis provides a foundation for future work to investigate time-varying and time-fixed state-level characteristics that may be responsible for differences among states in risk factors leading to each cause of death. Differences in smoking policy, Medicare coverage, access to care, segregation, and racism, for example, could all affect risk factors for ill health and vary in magnitude by state.

## Methods

### Description of the mortality data

We used software maintained by the National Cancer Institute’s Surveillance, Epidemiology, and End Results Program (SEERStat) to extract data on the number of deaths and the population size in all 50 US states and the District of Columbia between 1969 and 20 1 3.^28^ The mortality data were originally collected by the National Center for Health Statistics (NCHS) from death certificates and stored by the National Vital Statistics System.^29^ The population data were derived by the NCHS using intercensal bridged-race population estimates. We used data on the number of deaths and the population size according to race (black or white), gender (male or female), age group (<1 year old, 1–4 years old, followed by 5-year age groupings, with a final grouping of 85 or more years of age), year (1969 to 2013), and cause of death. Cause of death was originally recorded using the International Classification of Diseases (ICD) codes versions 8 (1969–1979), 9 (1979–1998), and 10 (1999–2013). These codes were aggregated into six broad causes of death for this analysis: Cancers, Cardiovascular disease, Communicable disease, Injuries, Non-communicable disease, and All other causes **(Table S1)**.

### Statistical challenges

The data structure produced two statistical challenges that were addressed by our modeling strategy. Firstly, because of confidentiality, NCHS suppresses counts of deaths between 1 and 9,^30^ that we needed to impute. Secondly, several states had small black populations which produced sparse counts in some state-year-race-sex-age-cause strata. In these cases, the estimation of the mortality rates is imprecise, and can produce abrupt changes in the estimated mortality rates over time that are unrealistic. To better estimate the underlying mortality rates, we used a method that produces smoother age-specific mortality trends.

### Estimation of smoothed mortality rates using a Bayesian time-series model

We used a Bayesian time-series model to smooth the death counts across years and estimate the underlying mortality rate. Specifically, we used a Bayesian generalized linear model with a Poisson likelihood to model the number of deaths in each state-year-sex-race-age-cause stratum as a function of the at-risk population, for strata with zero or more than 9 deaths. The mortality rates over time within each stratum were assigned an autoregressive prior, where the log mortality rate was assumed to be Gaussian centered at the previous year’s mortality rate. A non-informative gamma prior was used to estimate the inverse variance. For censored strata with between one and nine deaths, the deaths were modeled using a truncated Poisson likelihood, and otherwise the model remained the same. The truncated Poisson was bound between one and the larger of nine or the stratum’s population size, which could be less than nine.

Models were fit using Markov chain Monte Carlo (MCMC). These models were run separately for every combination of state, race, gender, and cause of death. They were fit using two chains of length 10,000, with a 2,000 iteration burn-in period to ensure convergence. One thousand iterations remained after burn-in and sampling every eighth iteration to reduce autocorrelation among the MCMC samples. Each iteration produced death count estimates that were smoothed over time for every cause of death within each age group, which were used to estimate life expectancy, the life expectancy gap, and perform a cause of death decomposition. These 1,000 decompositions form the posterior distribution from which the credible intervals were estimated using the 2.5^th^ and 97.5^th^ percentiles.

### Decomposition of the life expectancy gap by cause of death

To calculate life expectancy at birth, the smoothed cause-specific death counts were aggregated to estimate the total deaths in each age group, and standard life table methods were applied.^31,32^ Life expectancy at birth (in years) was calculated for blacks and whites in each state and year, by gender. The difference in life expectancy between blacks and whites was calculated by subtracting the black life expectancy from the white life expectancy in each state, year, and gender.

To examine the components of the difference in life expectancy between blacks and whites, we performed a decomposition analysis by cause of death using the Arriaga method.^33^ This method partitions the black-white difference in life expectancy by cause of death, which can be interpreted as the amount of the gap, in years, that can be attributed to higher cause-specific mortality rates among blacks. For example, a decomposition of a gap of five years may find that two of the five years is attributable to a higher rate of mortality due to cardiovascular disease among blacks compared to whites. If the decomposition produces a negative estimate this implies that whites have a higher rate of mortality for injuries compared to blacks. Thus, for each state, the contribution of each of the six causes to the black-white life expectancy gap was estimated for each year, producing 40 state-level trends and their credible intervals.

We also calculated state-level time trends in age-standardized cause-specific mortality rates separately for each combination of race and gender, to aid the interpretation of the decomposition results. We used the US standard population from the year 2000.^34^

### Description of the trend in each cause of death’s contribution to the gap and identification of divergent states

To describe the national trend in each causes’ contribution to the life expectancy gap over time, we used random effects meta regression to model each state’s mean contribution estimated from the previous step as a function of a state random effect and a linear spline for year (estimated using four internal knots). The model was inversely-weighted by the standard error of the mean estimate in each state-year. To identify states divergent from the national trend, we compared the state-specific trends estimated in the previous step to the national pattern estimated by setting the random effect to zero.

### Data Sharing

A replication data set including the raw data and statistical code to reproduce the manuscript will be made publicly available on github soon: https://github.com/corinne-riddell/BlackWhiteMortalityGap/.

## Acknowledgements

We extend our thanks to Dr. Richard Maclehose, who developed the initial version of the code to perform the Bayesian analysis as part of the research article “Trends in the Black-White Life Expectancy Gap Among Us States, 1990–2009”, by Harper S, MacLehose RF, and Kaufman JS.

Corinne Riddell is a post-doctoral researcher at McGill University, whose salary was supported by Dr. Kaufman’s Canada Research Chair in Health Disparities. Kathryn Morrison is a post-doctoral researcher at McGill University. Sam Harper was supported by a Chercheur Boursier Junior 2 from the Fonds de la Recherche en Santé du Québec. Jay Kaufman was supported by a Canada Research Chair in Health Disparities.

## Appendix

**Table S1:** Cause of death groupings

| Broad cause of death | Included causes of death |
| --- | --- |
| Cancers | Lip; Tongue; Salivary gland; Floor of mouth; Gum and other mouth; Nasopharynx; Tonsil; Oropharynx; Hypopharynx; Other oral cavity and pharynx; Esophagus; Stomach; Small intestine; Colon excluding rectum; Rectum and rectosigmoid junction; Anus, anal canal and anorectum; Liver; Intrahepatic bile duct; Gallbladder; Other biliary; Pancreas; Retroperitoneum; Peritoneum, Omentum and mesentery; Other digestive organs; Nose, nasal cavity and middle ear; Larynx; Lung and bronchus; Pleura; Trachea, Mediastinum and other respiratory organs; Bones and joints; Soft tissue including heart; Melanoma of the skin; Non-melanoma skin; Breast; Cervix uteri; Corpus uteri; Uterus, NOS; Ovary; Vagina; Vulva; Other female genital organs; Prostate; Testis; Penis; Other male genital organs; Urinary bladder; Kidney and renal pelvis; Ureter; Other urinary organs; Eye and orbit; Brain and other nervous system; Thyroid; Other endocrine including thymus; Hodgkin lymphoma; Myeloma; Acute lymphocytic leukemia; Chronic lymphocytic leukemia; Other lymphocytic leukemia; Acute myeloid leukemia; Acute monocytic leukemia; Chronic myeloid leukemia; Other myeloid/monocytic leukemia; Other acute leukemia; Aleukemic, Subleukemic and NOS; Miscellaneous malignant cancer. |
| Cardiovascular disease | Diseases of heart; Hypertension without heart disease; Cerebrovascular diseases; Atherosclerosis; Aortic aneurysm and dissection; Other diseases of arteries, arterioles, capillaries. |
| Communicable disease | Tuberculosis; Syphilis; HIV (1987+), Septicemia; Other infectious and parasitic diseases; Pneumonia and influenza |
| Non-communicable disease | Diabetes Mellitus; Alzheimers (ICD9 and 10 only); Chronic obstructive pulmonary disease and allied conditions; Stomach and duodenal ulcers; Chronic liver disease and cirrhosis; Nephritis, Nephrotic Syndrome and Nephrosis. |
| Injuries | Accidents and Adverse Effects; Suicide and Self-inflicted Injury; Homicide and Legal Intervention. |
| All other causes | Complications of pregnancy, childbirth, puerperium; Congenital anomalies; Certain conditions originating in perinatal period; Symptoms, signs, and ill-defined conditions; Other cause of death. |

## Figure Legends

Figure 1: The bands illustrate the 95% credible interval around the mean trend line. States with more imprecise estimates (often due to small black populations) have wider credible intervals. The estimated life expectancy gap in California in 1972 is much higher than in surrounding years. This anomaly is also illustrated in Figure 3, and is a product of an unexpectedly low CVD-related mortality rate among white males in California in 1972, likely denoting a coding error. Thus the data for this state and year should be interpreted with caution.

Figure 3: The blue line and light blue band illustrate the mean and 95% credible interval for each state’s estimated trend in the contribution of CVD to the black-white life expectancy gap over time. The red line is the same across the panels and illustrates the national pattern in the contribution, to add in the identification of similar and divergent states. The early spike in California’s trend line illustrates the unexpectedly high estimate of the CVD contribution to the black-white life expectancy gap among California males in 1972. This anomolous estimate reflects the unexpectedly low CVD-related mortality rate among white males in California in 1972, and likely denotes a coding error. Thus the data for this state and year should be interpreted with caution.

